# The microstructure investigation of plant architecture with X-ray microscopy

**DOI:** 10.1101/729533

**Authors:** Wenting Zhang, Tao Guo, Ke Chen, Ting La, Philipp Alexander Bastians, Chunjie Cao

**Author notes:** Wenting Zhang and Tao Guo contributed equally to this work.

## Abstract

**Background:** In recent years, the plant morphology has been well studied by multiple approaches at cellular and subcellular levels. Two-dimensional (2D) microscopy techniques offer imaging of plant structures on a wide range of magnifications for researchers. However, subcellular imaging is still challenging in plant tissues like roots and seeds.

**Results:** Here we use a three-dimensional (3D) imaging technology based on the ZEISS X-ray microscope (XRM) Versa and analyze several plant tissues from different plant species. The XRM provides new insights into plant structures using non-destructive imaging at high-resolution and high contrast. We also developed a workflow aiming to acquire accurate and high-quality images in the context of the whole specimen. Multiple plant samples including rice, tobacco, *Arabidopsis* and maize were used to display the differences of phenotypes, which indicates that the XRM is a powerful tool to investigate plant microstructure.

**Conclusions:** Our work provides a novel observation method to evaluate and quantify tissue specific differences for a range of plant species. This new tool is suitable for non-destructive seed observation and screening.

## Background

Since the invention and development of microscopes, it has extended human vision substantially. The observation of cellular and subcellular structures using microscopes have broadened our knowledge to understand the biological world more efficiently [1]. ZEISS and many other microscopy manufacturing companies are spending an enormous amount of time and resources developing higher resolution microscopy systems to assist scientists acquire more detailed images in their research fields. From the single cell organism blue-green algae (*Cyanobacteria*) to over a hundred-meter-tall giant tree (*Eucalyptus regnans*), plants display versatile morphologies to survive in different environments. Therefore, utilizing microscopy techniques to study the cellular and subcellular and physiological traits is essential in plant research.

In the 21^st^ century, microscopy companies provide a variety of measurement techniques for scientists. With the assistance of electron microscopy, plant scientists can observe the cell surface and the detailed structure of organelles, and even decipher the structure of proteins [2-4]. Optical microscopes, including upright and inverted microscopes, provide powerful solutions for cellular observations as well [5, 6]. Since the application of green fluorescent protein, confocal microscopy and various fluorescence related techniques advanced the biological research field [7]. Furthermore, the methods to analyze the corresponding data have been developed at a similar pace [8]. For larger sample observation, stereoscopic microscopy offers non-destructive and detailed insights to identify the tiny differences in between samples. Nowadays, plant scientists can observe nearly all kinds of samples with appropriate microscopes.

After the discovery of X-rays in 1895, X-ray imaging was rapidly used in biological science [9]. This unique technique has been further developed to analyze material composition, facilitate paleontological measurements, and detect components in metals, etc. [10]. Phytologists use x-ray computed tomography (microCT) to scan and observe plant materials non-destructively, especially in forestry morphometry [11]. High-resolution computed tomography (HRCT) is a well-established method to observe the plant vascular system in three dimensions [12-15]. This method was used to investigate and understand the formation of emboli in saplings’ xylem during cycles from drought to re-watering [16]. The plant root formation in the soil is another research focus to utilize HRCT imaging and analysis [17-19]. This method enables scientists to trace the development of roots and foresee the plant’s growth status. This can be used to select crop species to match the conditions of individual areas. Furthermore, leaves and seeds are imaged and characterized using HRCT [20-22].

ZEISS has developed high resolution and high contrast lab X-ray microscopy series since 2000. The ZEISS XRM uses an innovative two-stage magnification technology, which is different from traditional microCT. It maintains high-resolution with increasing sample size and obtains ultra-high contrast imaging with biological low-density materials. Our report suggests that XRM is an excellent tool to analyze cellular structures in plants. Furthermore, we demonstrate new ways to compare different plant phenotype samples, which will further increase our understanding of plant anatomy and function and expands the portfolio of 3D imaging methods. Moreover, we conducted several case studies using the XRM system for 3D observation, and our data suggest a bright future using XRM in the field of plant research.

## Methods

### Plant Materials and Conditions

The *Oryza sativa* wildtype FAZ1 and *gsn1* mutants were grown in fields close to Shanghai. The young spikelets and stems were obtained from three-month-old plants and fixed in FAA solution (50% ethanol, 5% glacial acetic acid, 5% formaldehyde). Mature seeds of tobacco, maize W22, rice FAZ1 and *gsn1* mutants were harvested from fields close to Shanghai under natural desiccated conditions. The rice *indica* variety “TeQing” (TQ) seedlings were grown in an artificial environment at 25°C for 5 days. The plants were grown in glass tubes in solid 1/2 Murashige and Skoog (MS) medium. For *Arabidopsis thaliana*, the ecotypes of Col-0 and Sij-4 plants were grown in soil under long days condition (16-hr-light/8-hr-dark cycles) at 21°C in phytotron.

### Plant pretreatment

There are three conditions of plant material used in this research. The first condition was naturally desiccated plant material which was the state of the tobacco, *Arabidopsis*, and maize and rice seed samples. The second condition is fresh plant material which was directly used for XRM observation like the fresh rice seedlings. The last condition was fixed and dehydrated plant material. The plant material was incubated in formaldehyde-acetic acid alcohol (FAA) overnight at 4°C. The infiltration of FAA was assisted by exposing the immersed plant material to vacuum, the target pressure is near 0.2 bar for 10s. Repeat this step until all the plant materials sinking down the FAA. Next, the plant material was dehydrated in graded ethanol (5 min at 50%, 5 min at 75%, 5 min at 85%, 5 min at 95% and 5 min at pure ethanol). Then, the samples were desiccated in an automated Critical Point Dryer (Leica EM CPD 300).

### XRM observations

The specimens were mounted with epoxy resin and scanned with a ZEISS Xradia 520 Versa X-ray Microscope (Carl ZEISS (Shanghai) Co. Ltd.). Specimens were mounted on the holder with a centrifuge tube or Kapton® tube as adapter and rotated horizontally by 360° or 180° + fan, pausing at discrete angles to collect 2D projection images, which were then computationally combined to produce a 3D reconstruction of the specimen’s volume dataset. For plant sample, usually, we use low kV for imaging, scanning energy is 40kV/3W – 50kV/4W. All parameters of scanning by XRM are recorded in Additional file 1: Table S1. At relatively low voltage, low density plant specimens can produce enhanced phase contrast effect. Voxel sizes vary from 0.3 to 5.0 μm depending on the field of view.

### Image processing and 3D remodeling

Image stacks of about 1000 virtual slices with 0.3-5.0 μm voxel sizes were acquired on XRM. XRM3DViewer and Dragonfly software were employed for 3D image processing, including image 3D reconstruction, segmentation and rendering. The interior structure of the XRM images were adjusted using gray-scale based threshold selections. The data segmentation can be focused on quantitative analysis of plant structure, such as porosity, wall thickness, cell number, aspect ratio, surface area and others. Consecutively, this data can be used to characterize plant growth traits.

## Results

### Architecture of Xradia 520 Versa X-ray Microscopy

In our research, we used the ZEISS Xradia 520 Versa X-ray Microscope to observe multiple different plant species, including rice (*Oryza sativa*), maize (*Zea mays*), tobacco (*Nicotiana benthamiana*) and *Arabidopsis* (*Arabidopsis thaliana*) (Fig. 1a). Since the development of the Xradia 520 Versa XRM in 2013, the high-resolution and outstanding contrast make the XRM applicable to a wide-range of applications in the mining industry, the electronic industry, and the material and life science research. The ZEISS Xradia 520 Versa XRM contains a source emitting X-rays which directly transmit through the immobile sample. Then, the transmitted X-rays hit a scintillator which converts the signals into visible light. At the back of the optical magnification unit, a large detector receives the visible light signals and an image is generated (Fig. 1b). The center unit of the XRM is composed of three pieces: the source that emits X-rays, the sample holder that allows X, Y and Z movement and rotation, and the detector that receives the transmitted signals (Fig. 1c). The samples are mounted on the aluminum holders. Then, the samples are rotated, and paused at discrete angles to collect 2D projection images. Subsequently, the projection images are combined computationally yielding a 3D reconstruction of the sample’s volume. It takes tens of minutes to several hours to scan different samples at a resolution of several micrometers to sub-micrometer, respectively. During the scanning, the samples must be stable and stationary. The challenge of fresh plant samples is the water evaporation, which causes scanning failures due to the sample deformation. In addition, due to the similar x-ray absorption rate of biological tissue and water, fresh and hydrated plants cannot achieve high resolution and sufficient contrast. Fresh leaves, roots, stems, seeds and other plant parts were tested. For natural dehydrated plant samples, such as seeds, we performed the scanning without special sample treatments. Fresh hydrated samples were sealed with Para-film or mounted into a Kapton® tube before the XRM scan. We recommend critical point drying for samples where the features of interest need high resolution and therefore long scanning time, such as rice flowers, *Arabidopsis* stems. We applied this pretreatment to plant culms, young spikelets, and *Arabidopsis* flowers and legumes and obtained high contrast high-resolution 3D data.

**Fig. 1.**
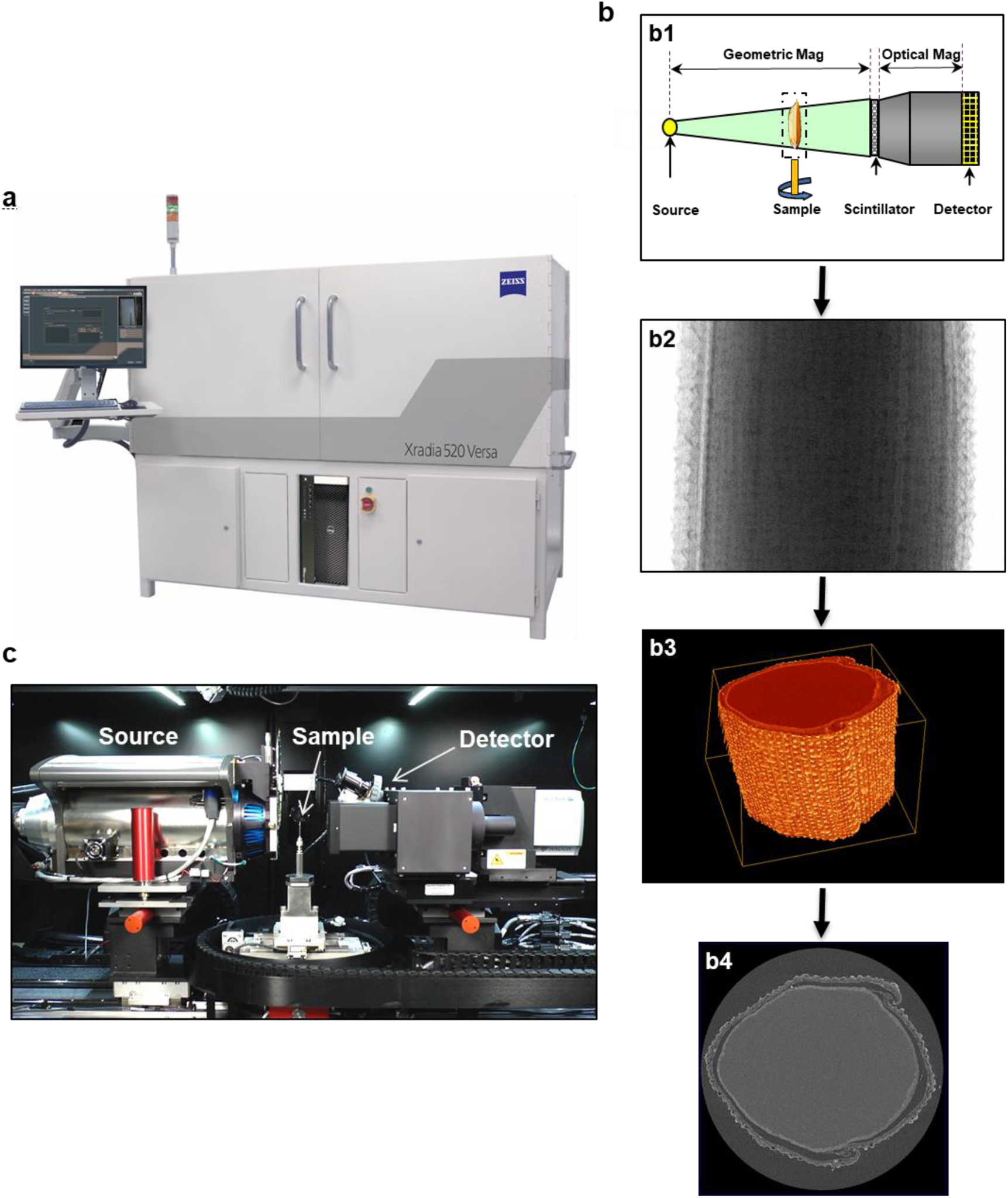
A brief introduction of the Xradia 520 Versa XRM. **a** A picture of the Xradia 520 Versa XRM. It contains an integrated working station and an operating system which is convenient for users to do the sample alignment and define the scan setting. **b** A simple workflow on the Xradia 520 Versa XRM. The X-rays are spatially mapped on the scintillator in front of the objectives. The scintillator converts X-ray into visible light, which is further magnified by the objectives. The sample is rotated, and paused at discrete angles to collect 2D projection images. Projection images are computationally combined to generate a 3D reconstruction of the sample’s volume. Researchers can investigate the results by roaming through the virtual slices. **c** The center unit of the Xradia 520 Versa XRM system.

### The XRM observation of the rice root

The essential requirements of a plant, such as the water absorption, the nutrient availability and eventually the plant development are determined by root structures [23]. Therefore, we evaluated fresh rice root samples by XRM first. The samples were grown for 7 days after germination under normal planting conditions, then the roots were cut to a length of 3 cm. The sample was sealed within a Kapton®tube before the scan as shown (Fig. 2a). After scanning, the 2D projections were computationally reconstructed into a 3D dataset. The ZEISS XRM3DViewer Software was used to render and visualize the 3D volume (Fig. 2b). The 3D data can also be visualized and investigated by three 2D projections, which are orthogonally oriented to each other, representing spatial interpolations of the 3D volume’s intensity values. Each scanning iteration generates about 1,000 images of 2D projections from specific angles. These 2D projections can be also analyzed by the ZEISS XRM3DViewer software. Although the data reflects the true shape and intact internal structure of the hydrated root samples, the intrinsic contrast is limited by the differential absorption rate of the plants’ structural composition. In addition, the superimposed structures do not allow for a clear detection of the boundaries of individual features. In order to display the general cell morphology, we chose four reconstructed 2D projections to observe the internal structure of rice roots as examples (Fig. 2c-2f). The video was rendered to display all the reconstructed 2D projections of all three cardinal axes to provide an overview of the acquired data (Additional file 2: Video S1). Using the reconstructed 2D projections, we observed the primary root region and identified epidermis cells, vascular bundles and the Casparian band. Next, we investigated the cellular arrangement of the root (Additional file 3: Fig. S1). We found that the lateral root cells are smaller than the main root cells (Additional file 3: Fig.S1A and S1C). The young roots contained small and compact xylem cells which were surrounded by large and loose sieve cells. Due to the missing silicon in the growth medium, the Casparian cell band is thinner which has been reported before [24]. Although the fresh sample contained a high moisture content which reduced contrast in 2D projections, the boundaries of xylem and phloem were still visible. This suggests that the XRM is an excellent tool to display the cellular structure of plant root tissue without any treatment.

**Fig. 2.**
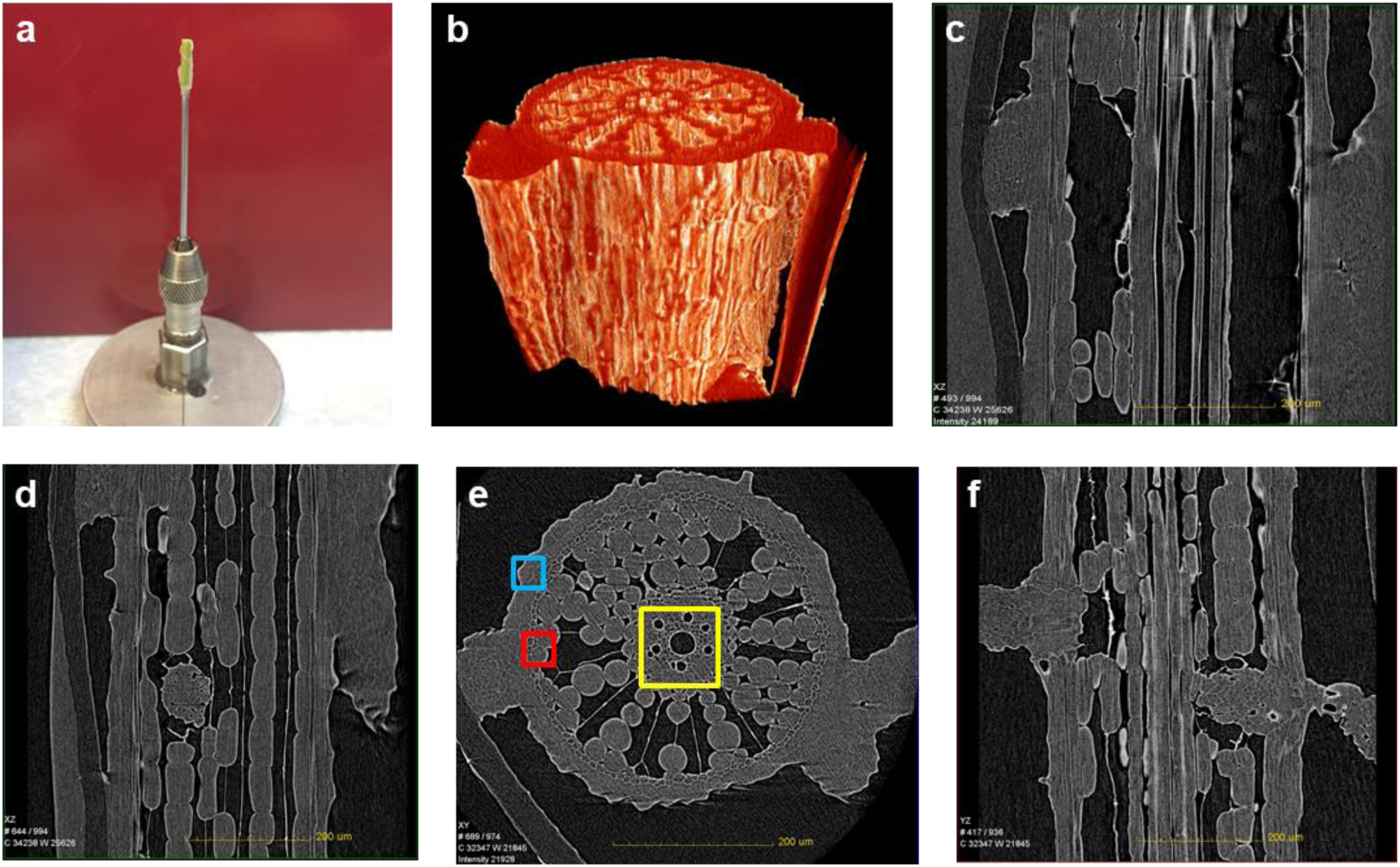
The root’s microstructure image of the rice *indica* variety TeQing (TQ). **a** Each fresh rice root sample is mounted in a Kapton® tube and the tube is mounted on an aluminum holder. **b** The rendered 3D data is colored to represent voxel intensities. **c-f** reconstructed 2D projections of (**b**) in different orientations. Scale bars, 200 μm. The dark area indicates the aerate tissue of the xylems in the rice root. The grey shows the parenchymal cells of xylems. All the scanning parameter are listed in the additional file 1.

### The XRM observation of the rice stem

Next, we investigated samples from the rice stem using XRM. The plant samples were pretreated to increase the contrast and quality of the 2D projections. The stem samples were pretreated using critical point drying (CPD). We applied a CPD protocol which is conventionally used for sample imaging in scanning electron microscopy as described here [25] (Additional file 4: Fig. S2A). After scanning and image processing, the XRM3DViewer visualized the shape of the stem (Additional file 4: Fig. S2B). Next, we investigated the vascular bundles and the cell sizes of the stem tissue using appropriately oriented 2D projections (Additional file 4: Fig. S2C). The scanning parameters yielded a voxel size of 2.1 μm and the imaging results reflect the structures of the stem sufficiently, e.g. the inner longitudinal surface; however, the images cannot resolve detailed microstructures like parenchymal cells of stems. The reconstructed 2D projections show the cross sections of the cell walls prominently as a bright circle which is substantially improved due to the critical point drying (Additional file 4: Fig. S2D-S2E). Therefore, our images from the 2.1 μm scanning suggests that the XRM can reflect the general structure of the rice stem from the low-resolution scan. If higher resolution images are required, the samples must be trimmed or cut into smaller pieces. We cut the stem tissues into a smaller piece and mounted it onto the sample holder to perform high-resolution scanning (Fig. 3a). The field of view of the stem sample contained an intact vascular bundle and a fraction of the parenchymatous tissue (Fig. 3b). The images in Fig. 3c-3f clearly show the cell structure and arrangement in the rice stem. As expected, the XRM resolved the cell walls clearly. The more superficial portion of the rice stem is densely packed with cells, and gradually becomes more dispersed towards the center. Next, we used the measurement tools of the ZEISS XRM3DViewer software to evaluate the average cell size (Fig. 3g). These results suggest that the XRM is capable of cell counting, cell size measurement and evaluation of the cellular organization.

**Fig. 3.**
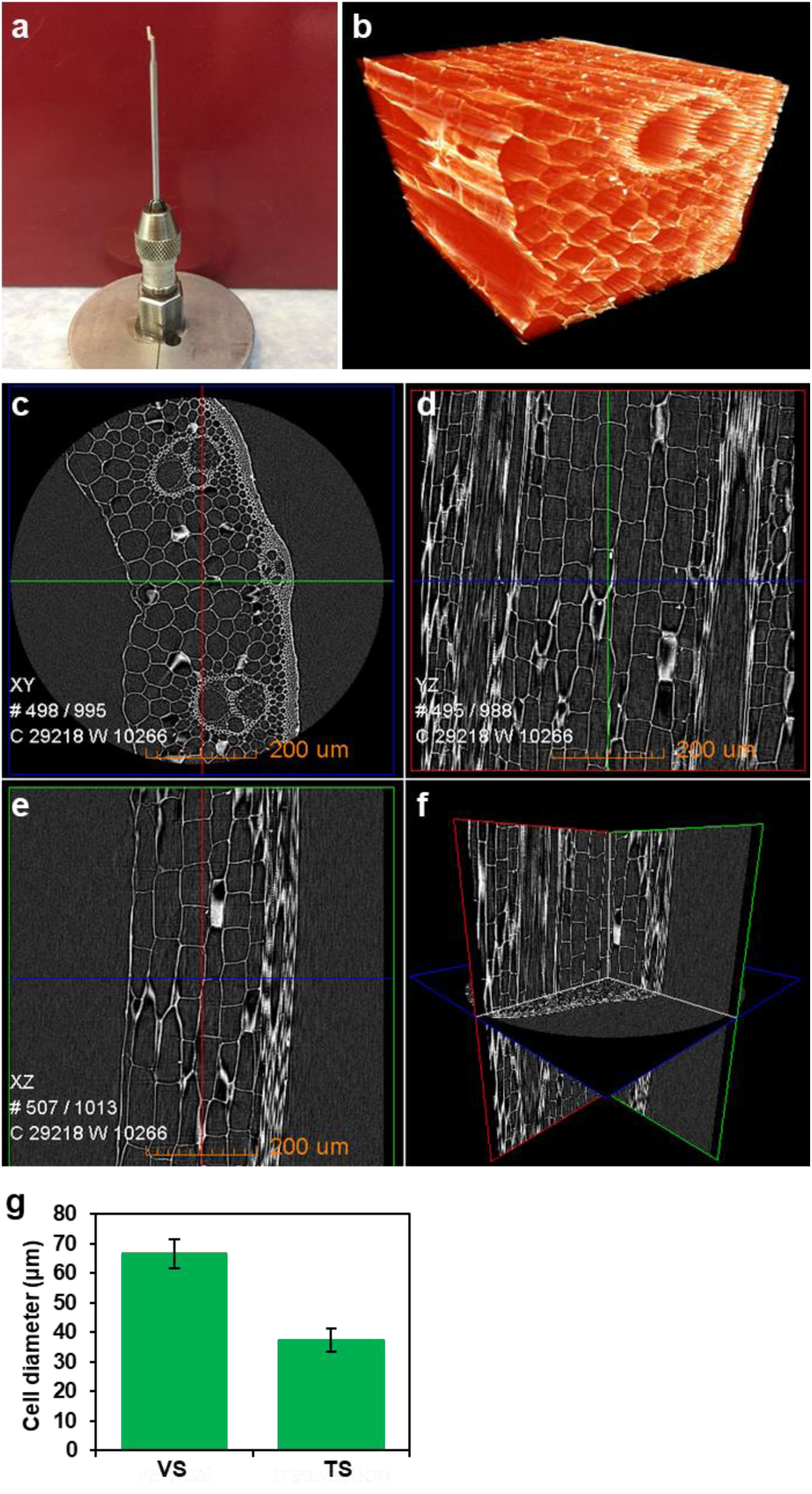
The stem microstructure of the *indica* rice variety TQ. **a** The critically point dried rice stem sample was loaded on the aluminum sample holder before the XRM scan. **b** The rendered 3D data is colored to represent voxel intensities. **c** The reconstructed 2D projection shows the cross-section of the rice stem rim. The white circles represent the cell walls. **d**-**e** the images show the orthogonally oriented reconstructed 2D projections indicated by the red and green line in **c**, respectively. The cell walls appear as stretched boxes which are oriented parallelly to the stem axis. **f** The blue, red and green viewports are oriented orthogonally to each other. Each viewport shows corresponding 2D projection. Scale bars, 200 μm. **g** The statistical analysis of stem cell diameter (n=50). VS, vertical surface; TS, transection surface. The error bar indicates SEM.

### The XRM observation of the rice grain

Next, we applied the XRM to mature grains of rice (Fig. 4a). We mounted the grain from the *indica* rice FAZ1 (Fengaizhan-1) on the sample holder and scanned for 1.1 hours to yield a 3D overview image yielding a voxel size of 9.3 μm. Next, we scanned twice for 1.8 hours to image the areas indicated (Fig. 4b-4d). The rendered 3D volume shows clear the distribution of the testa, which can also be observed using a stereo microscope (Fig. 4a, c). The reconstructed 2D projection parallelly to the long axis of the seed shows clear withered anther in region I and the embryo in region II (Fig. 4d). Two reconstructed 2D projections parallel to the axis of the seed grain were displayed (Additional file 5: Fig. S3), which shows a detailed morphological pattern of the starch arrangement. These two regions were further investigated by acquiring another 3D image at higher magnification for 1.8 hours (Fig. 4e, f). Two reconstructed 2D projections were selected for each dataset to evaluate the cellular structure of the endosperm and embryo (Fig. 4g-j). We found that in the mature part of the grain, the endosperm is filled with non-cellular structures, such as starch (Fig. 4i, j), and the embryo is composed of densely arranged cells (Fig. 4h). Next, we imaged the joint area of the inner and outer grain glume at a higher magnification for 3.5 hours yielding a voxel size of 0.7 μm (Fig. 4k-p). The 3D reconstruction of the joint area shows clearly how cells arranged in the FAZ1 rice grain (Fig. 4l, m, o and p). The imaging results indicate that the rice grain is a suitable specimen for XRM scanning. The internal microstructure is resolved clearly and due to its non-destructive nature, the specimen can further be investigated using other imaging and analysis techniques.

**Fig. 4.**
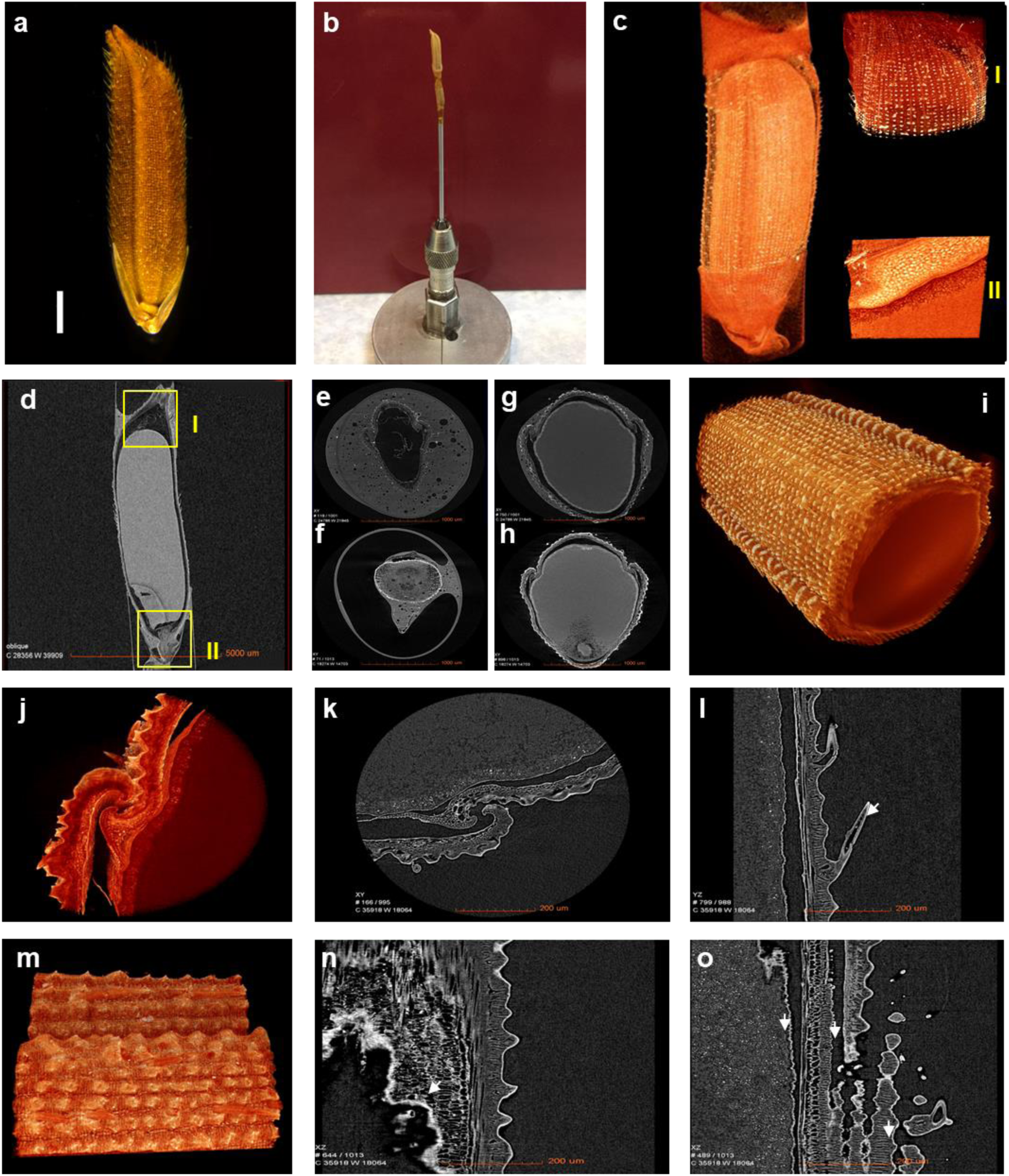
The grain microstructure image of the mature rice *indica* variety FAZ1. **a** Grain morphology of FAZ1. Scale bars, 1 mm. **b** The loaded holder with the dehydrated rice grain. **c** the rendered 3D data of the rice grain in (**b**) is colored to represent voxel intensities with a voxel size of 9.3 μm. The indicated area I and area II are the top and bottom of the rice grain and were scanned yielding a voxel size of 2.3 μm. **d** the reconstructed 2D projection of longitudinal section shows the internal morphology of the rice grain. The rectangles indicate the location of area I and II. **e**-**h** reconstructed 2D projection images of the top and bottom of area I and II (**d**). (**i**) 3D reconstruction with colored representation of the voxel intensities of the rice grain middle part. The reconstructed 3D images of the rice grain testa (**j, m**) and 2 different orientation of reconstructed 2D projections (**k, l, n, o**). All scale bars indicate the same length, 200 μm.

Besides the stem samples, we also scanned several critical point dried young spikelet specimens with the XRM system. We illustrate the microstructure investigation by using a comparative experiment design between the FAZ1 wildtype and the *gsn1* mutant specimen [25]. The *gsn1* rice mutant shows bigger grain size and less spikelets. The reconstructed 2D projection containing the longitudinal axis of the young spikelet reflected the number and the size of the primordia (Additional file 6: Fig. S4A and S4C). The FAZ1 sample showed more primordia that were smaller in size, which is consistent with the results of other studies [25]. Next, we scanned the glume of *gsn1* and FAZ1 and compared these regions (Additional file 6: Fig. S4B and S4D). We found that the size of cells located in the superficial portion of the FAZ1 glume is smaller in comparison to the *gsn1* glume, suggesting a faster cell division in *gsn1* (Additional file 6: Fig. S4B and S4D) [25]. These results suggest that the ZEISS XRM system is a suitable novel tool to observe the shoot apical meristem (SAM) and young panicles, which is a common analysis used by developmental botanists. In summary, the desiccative samples prepared with CPD is a suitable pretreatment method to display cellular morphology in plant tissue. Compared with the fresh samples, the CPD samples provide improved contrast in the reconstructed images obtained from the XRM observation showing detailed microstructures.

### The observation of maize seed by XRM

Next, we applied XRM to maize seeds. As a globally important cereal crop, an optimal and consistent production of maize is a major objective of farmers. Applying XRM to maize seed screening process can rapidly identify the shape and seed status without tedious dissection.

First, we attached the dehydrated maize seed at the top of the sample holder (Fig. 5a). Next, we scanned the maize seed at high resolution for 3 hours yielding a voxel size of 0.7 μm to resolve sufficient details for an internal microstructure analysis (Fig. 5b). The reconstructed 2D projections provided clear images of the endosperm, embryo and episperm (Fig. 5c). The endosperm is composed of two parts, the vitreous portion of the endosperm consists of more X-ray absorbing peripheral tissue yielding a brighter or higher signal intensity in the image. The floury endosperm showed a darker or lower signal intensity due to a composition that absorbs less X-rays. This brightness or signal difference was visible in the interior of the endosperm. Next, we scanned the vitreous and the floury portion of the endosperm for 6.5 hours yielding a voxel size of 0.3 μm. The brighter area is composed of a tight starch structure. Due to the dense packing of the starch granules, the reconstructed 2D projections were hardly showing any isolate particles (Fig. 5d, Fig. 5g-j). The vitreous portion of the endosperm is composed of a looser starch structure and the dark areas which contain water-soluble short chain carbohydrates (Fig. 5e, Fig 5k-n). In summary, we successfully used the XRM to observe the inner structure of the maize seed and distinguished the two components composing the endosperm. We propose that the XRM system is suitable as a powerful tool to screen seeds of mutants that alter the seed composition without performing cross-sections.

**Fig. 5.**
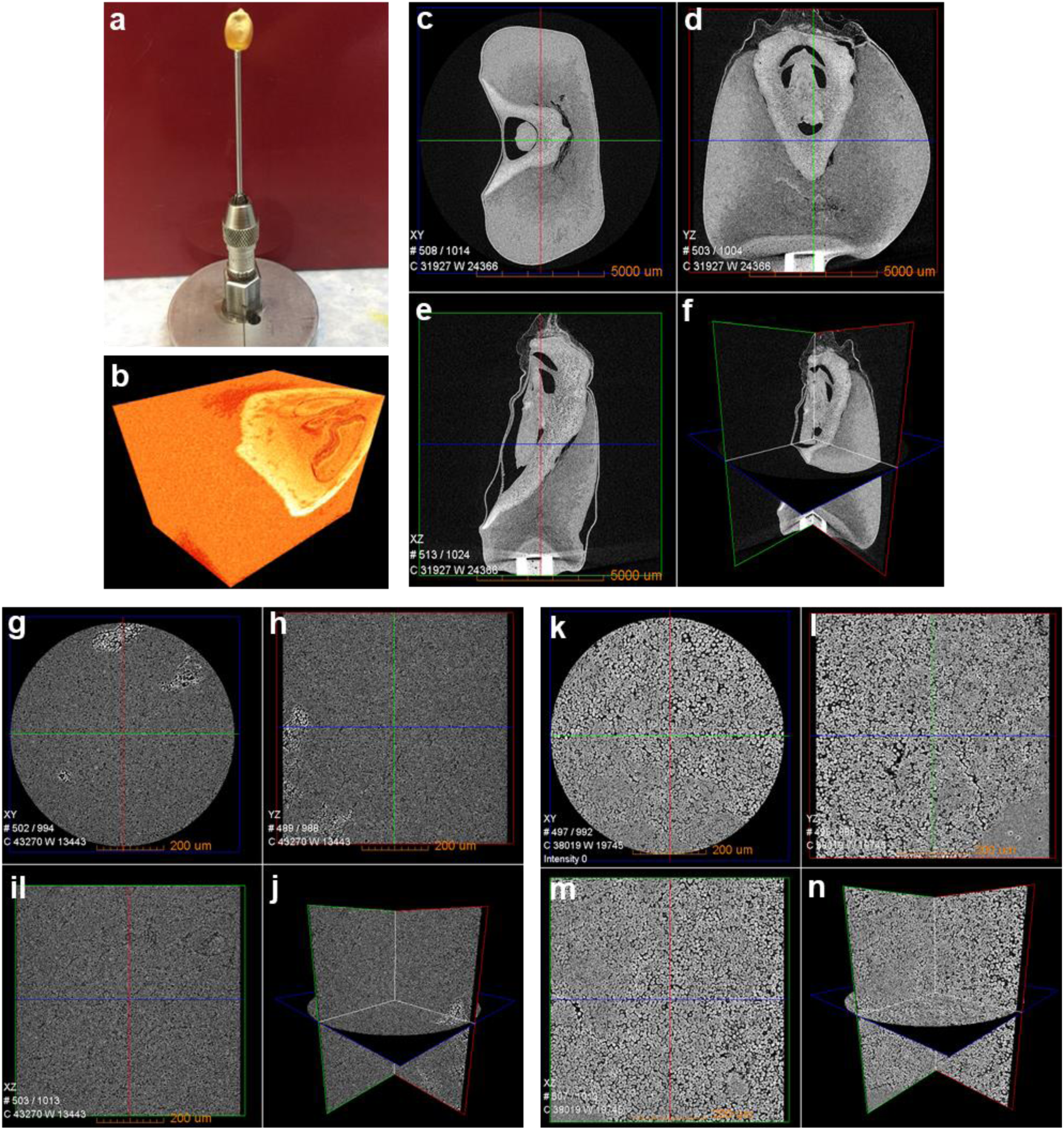
The microstructure XRM image of the natural dry seed of *Zea mays*. **a** a digital photography of the holder mounted with the naturally dehydrated maize seed. **b** the rendered 3D data of the maize seed in (**b**) is colored to represent voxel intensities with a voxel size of 9.5 μm. **c**-**f** the reconstructed 2D projection shows the cross-section of the maize seed orthogonal to the longitudinal seed axis. **d**-**e** the images show the orthogonally oriented reconstructed 2D projections indicated by the red and green line in **c**, respectively. (**f**) the images of **c-e** are combined and reflect the orthogonal orientation. Scale bars, 5000 μm. **g**-**j** the visualization of the reconstructed 2D projections of the vitreous endosperm with a voxel size of 0.7 μm. The starch granules are barely visible due to the dense packing. Scale bars, 200 μm. **k**-**n** the same visualization as in **c-f** and **g-j** showing the XRM scan result of the floury endosperm at a voxel size of 0.7 μm. Single starch granules can be differentiated. Scale bars, 200 μm.

### The observation of tobacco and *Arabidopsis* by XRM

Due to the small size of tobacco and *Arabidopsis* seeds, it is difficult to focuson their microstructures. We used mature tobacco seeds and obtained a reconstructed 3D XRM image (Fig. 6a). We scanned the seed for 6 hours and yielded a voxel size of 0.5 μm. The images clearly showed the boundaries of the cotyledon and the embryo area. A encapsulating sheath of sclerenchyma cells is protecting the seed. The reconstructed XRM data also resolved parenchymal cells and starch granules. We also scanned the *Arabidopsis* seeds from ecotypes Col-0 (Columbia-0) (Fig. 6e-h) and Sij-4 (Sidzhak-4) (Fig. 6i-l). The appearance of the Col-0 seed shows similarity to the Sij-4 seed with winkles around the seed (Fig. 6e+i). By choosing the appropriate reconstructed 2D projections one can easily identify the developing embryo, growing root, the polar of the seed, and two cotyledons linked with radicle. within direct comparison with the Sij-4 sample, we found less starch in the Col-0 sample. Considering that the Sij-4 is an ecotype of *Arabidopsis* grown in Uzbekistan [26], we inferred that the Sij-4 plants has a higher demand for energy stored as starch to survive due to the relatively low temperature environment.

**Fig. 6.**
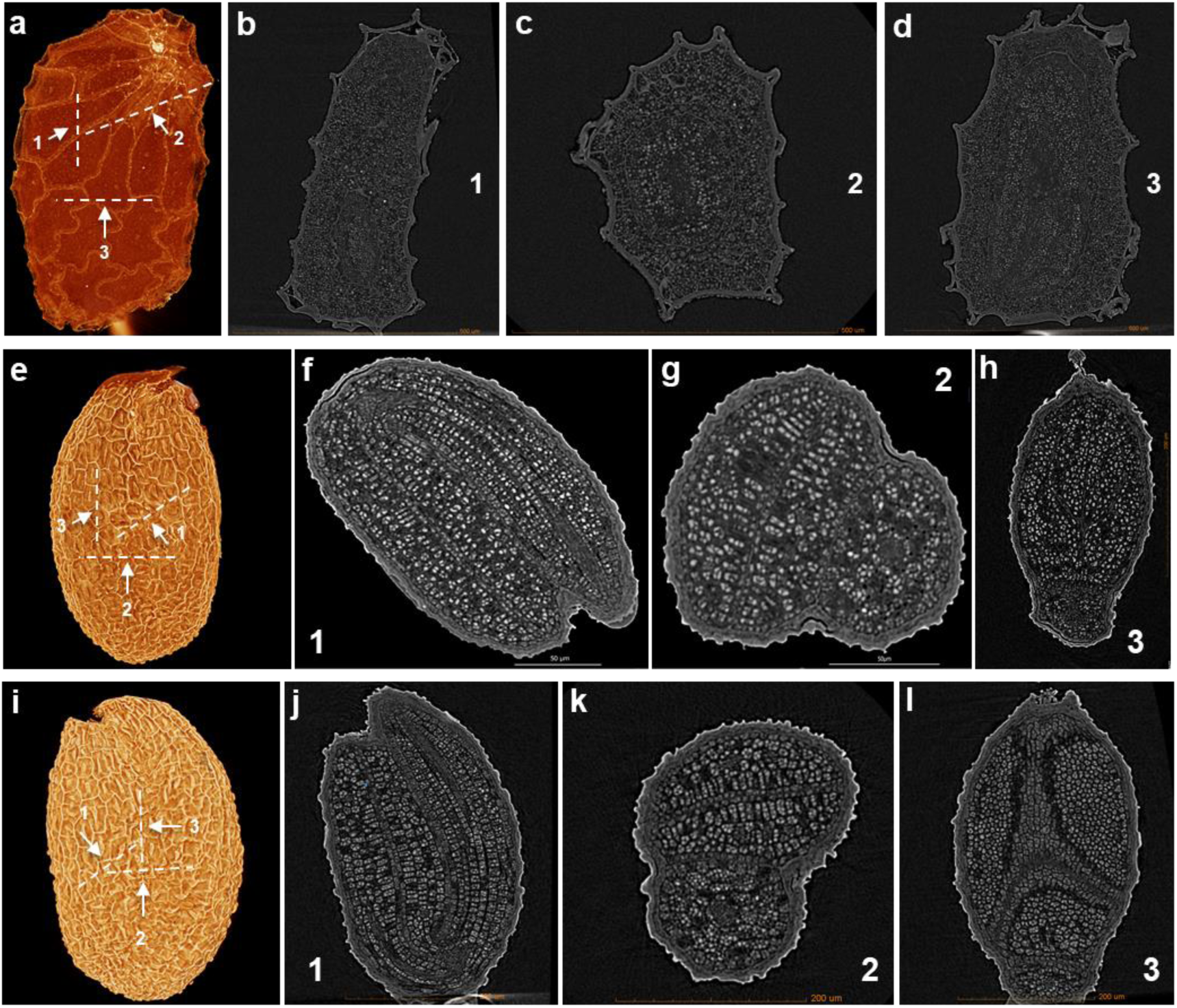
The microstructure XRM image of the tobacco and *Arabidopsis* natural dry seed. **a**-**d** the reconstructed and rendered 3D XRM data of the maize seed is colored to represent voxel intensities with a voxel size of 0.5 μm. Scale bars, 200 μm. **e**-**h** The same visualization as in **a-d** showing the XRM scanning results for the *Arabidopsis* Col-0 seed sample at a voxel size of 0.5 μm. The cotyledons range, the radicle area, the root growing spot and the embryo growing spot could be seen. Scale bars, 200 μm. **i**-**l** the same visualization as in **a-d** showing the XRM scanning results for the *Arabidopsis* Sij-4 seed sample at a voxel size of 0.5 μm. The cotyledons range, the radicle area, the root growing spot and the embryo growing spot could be seen. Scale bars, 200 μm.

We further analyzed the number and size of the cells. We obtained the 2D reconstructed image, then we segmented cells of the Col-0 seed and used the Dragonfly software to calculate the number of cells and their volume. The image processing software Dragonfly was used to divide the internal seed starch granules in the both *Arabidopsis* ecotypes (Additional file 7: Fig. S5A). The starch cell volume analysis showed that the Col-0 cell volumes range from 2.5 μm^3^ to 60 μm^3^. This histogram can be used as a fingerprint analysis to evaluate the quality of seeds and the development state(Additional file 7: Fig. S5B). Moreover, coloring the seed to reflect concavity and convexity provide additional information about the starch cell arrangements (Additional file 7: Fig. S5C and S5D) compared to the reconstructed 2D projections. These observations suggest that XRM is capable to observe the plant seed traits in a novel approach.

Next, we used XRM to observe the microstructures of *Arabidopsis* flowers and immature legumes (Additional file 8: Fig. S6). The structures of spores and stamens have been resolved (Additional file 8: Fig. S6A). High contrast spores is consisted of starches (Additional file 8: Fig. S6A). The image processing software XRM3DViewer was used to portray the pistils inside an immature flower bud with purple pseudo colors clearly discovering the pistil arrangement non-destructively (Additional file 8: Fig. S6B). We used Col-0 legumes to observe the fertilized immature seeds. Each seed contains thickened testa and a heart-shaped embryo (Additional file 8: Fig. S6C and S6D). To summarize this work, our data indicates that the ZEISS XRM system together with the Dragonfly software, provides a novel and efficient observation of a broad range of plant tissues.

## DISCUSSION

The field of plant research has been growing strongly and with it the demand for more accurate imaging methods to observe plant tissues is growing too. In recent years, the application of X-ray Microscopy in the Life Sciences is becoming an established imaging technique. This technology has been applied in insect taxonomy, paleontology and other Life Science focused research fields [27, 28].. However, limited exploration on how to apply this novel technology into plants observation was reported.

Recently, the X-ray Fluorescence Microscopy (XFM) technology has been widely used in elemental studies in plant science [29]. This new technology can monitor the distribution and concentration of elements in plant tissues. However, the resolution of XFM is limited, the resolution is similar to stereo microscope. Combining the XRM with XFM and electron microscopy can provide researcher a multi-facet observation of plant tissue containing element analysis and localization of metal tagged proteins. Various studies focused on using low resolution XRM to observe xylem structures in recent years [12, 13, 16, 30], indicating potential applications of XRM in plant science research. For researchers who are working on tree saplings, large samples are tough to use XRM to monitor their status in field, and only low resolution microCT is applicable. Such samples need to be cut in smaller pieces for scanning by the novel XRM system. Moreover, maize seeds have also been imaged using non-destructive density area calibration by microCT [20] as well as plant leaves [21, 31]. Engineers at ZEISS are working on developing new materials that can aggregate the X-ray, which will greatly expand the application of XRM into sub-micrometer resolution. The eventual goal is to solve the limitation of the sample size. This perspective makes XRM a promising tool for the Life Sciences. Our results show that the ZEISS X-ray Xradia 520 Versa microscopy can provide sufficient high-resolution to identify microstructure of most plant tissues. This technique provides faster and more efficient workflows for plant breeding and help researchers to identify and isolate target genes resulting in desired traits.

Here, we reported several applications based on the ZEISS Xradia 520 Versa X-ray Microscopy system focused on plant tissue observations. The presented XRM imaging workflow starts at the sample preparation for consecutive XRM imaging, highlights the results of the reconstructed data and indicates methods for data analysis applicable to plant phenotypic analyses (Additional file 9: Fig. S7). The XRM is a novel and powerful tool to study the internal structure of plant tissues that are precious or difficult to dissect. It also provides valuable information about the superficial surface and longitudinal surface which the cross-section is complicated to study until now. The software affiliated to the XRM system provides a convenient platform for segmentation of structures like cells and furthermore offers tools to do statistical analysis. Our data showed that the best contrast of the reconstructed data is yielded by applying a desiccation pre-treatment, followed by naturally dehydrated samples and fresh samples. However, even the XRM imaging result of fresh plant samples can be used to measure the cell size and numbers. Our work demonstrates potential applications for plant tissue measurements. There are still challenges for plant tissue observations, such as the limitation of resolution, the high cost of instruments etc. We hope this case study of XRM application will expand the tools in the plant science research.

## Acknowledgements

We thank Professor Hong-Xuan Lin (Shanghai Institute of Plant Physiology & Ecology) for sample supports. We thank Professor Xiaoshu Gao (Shanghai Ninth People’s Hospital) and Zhenhuan Xue (Carl Zeiss) on expertise and advice.

## Authors’ contributions

C.C. conceived and supervised the project, and C.C, W.Z. and T.G. designed the experiments. W.Z. and T.G. performed most of the experiments. K.C., T.L. and P.B performed some of the experiments. W.Z., T.G. and C.C analyzed data and wrote the manuscript. All authors read and approved the final manuscript.

## Ethics approval and consent to participate

Not applicable.

## Consent for publication

Not applicable.

## Additional Files

**Table S1**. The summary of scanning parameters of all samples used in this study.

**Video S1**. Combination of 2D projections in rice roots.

**Figure S1**. Detailed 2D projections of rice root in Fig. 1. (**A**-**B**) 2D projection intersecting the surface of the rice root with 0.7 μm voxel size. Scale bars, 200 μm and 100 μm respectively. Yellow square frame indicates the inset. (**C**-**D**) 2D projection of the lateral surface of the rice root with 0.7 μm voxel size. Scale bars, 200 μm and 100 μm respectively. Individual cell size is measured by blue lines and numbers.

**Figure S2**. The 2.1 μm voxel size scan of the rice stem tissues. (**A**) The loading platform with the mounted rice stem tissue. (**B**) The 3D reconstruction of the sample in (**A**) with computationally rendered. (**C**) 2D projections of GC young stem oriented in three orthogonal directions: the intersecting, longitudinal and lateral cut. Scale bars, 1000 μm. (**D**-**E**) 2D projections intersecting the surface of the stem. Scale bars, 1000 μm and 500 μm respectively.

**Figure S3**. Detailed insights into a mature rice grain. (**A**) 3D reconstruction of the rice glume scan at a voxel size of 2.3 μm. The white arrow indicates the glume hair. (**B**) 3D reconstruction of the rice endosperm scan at a voxel size of 2.3 μm. The white arrow indicates the glume hair. The white arrow indicates the starch granules. (**C**-**D**) 2D projection of a longitudinal cut of the top seed with 9.5 μm voxel size. Scale bars, 1000 μm and 500 μm respectively. The yellow square frame indicates the inset. The blue measurements in (**D**) show the different cell widths. (**E**) 3D reconstruction of the rice embryo scan at a voxel size of 2.3 μm. (**F**) The 3D rendering shows the same data as in (**D**) from a more superficial region. (**G**-**H**) 2D projections of a longitudinal cut of the seed embryo with a voxel size of 9.5 μm. Scale bars, 1000 μm and 500 μm respectively. The numbers on (**D**) shows the different tissues width.

**Figure S4**. Young spikelet and glume comparison of FAZ1 and *gsn1*. (**A**) 2D projection of a longitudinal cut of FAZ1 and *gsn1* young spikelet. Scale bars, 500 μm. The FAZ1 and *gsn1* were scanned with a voxel size of 1.6 μm and 0.9 μm respectively. (**B**) 2D projections intersecting the glumes of FAZ1 and *gsn1*. Scale bars, 200 μm. The FAZ1 and *gsn1* were scanned at a voxel size of 0.7 μm and 0.5 μm respectively. 2D projections of a FAZ1 (**C**) and *gsn1* (**D**) young spikelet in 3 orthogonal orientations: the intersecting, longitudinal and lateral cut. Scale bars, 200 μm. (**D**) 2D projections of young spikelet in 3 different direction. Each direction represents the intersecting cut, longitudinal cut and lateral cut. Scale bars, 500 μm.

**Figure S5**. Segmentation and rendering of *Arabidopsis* Col-0 seed with the Dragonfly software. (**A**) The 3D rendering of the Col-0 seed after XRM scanning and data segmentation using the Dragonfly software. Each cell is divided into a separate 3D segmentation and rendered according to the volume. (**B**) A normalized histogram of the cell volume of the Col-0 seeds. The column colors are consistent with (**A**). (**C**-**D**) Two Col-0 seed 3D renderings color mapped by concavity and convexity (**C**) and flat embellishment (**D**).

**Figure S6**. The observation of *Arabidopsis* flower buds and legumen. (**A**) 2D projection from a longitudinal cut through the dehydrated Col-0 flower bud. Scale bars, 500 μm. The voxel size of the flower bud scan is 0.9 μm. (**B**) The fake color embellished 3D rendering of the *Arabidopsis* flower bud. Scale bars, 100 μm. (**C**) 2D projection from a longitudinal cut through the Col-0 legumen. Scale bars, 100 μm. (**D**) The pseudo color embellished image of *Arabidopsis* legumen. Scale bars, 500 μm.

**Figure S7**. The imaging workflow for XRM to observe plant samples. Most plant samples belong to three different categories, including fresh samples, dehydrated samples and desiccated samples. We used CPD method to creat dehydrated samples. After preparation, the samples are loaded in the XRM system. Then, the scanning parameters for each sample are defined and the scan is initiated. During the scanning procedure, the sample should remain still. The scan time depends on the resolution and the x-ray opacity of the sample. The ZEISS system provides powerful and convenient image processing tools to reconstruct the 2D projections.

