## Supplemental Figures for "The microstructure investigation of plant architecture with X-ray microscopy"

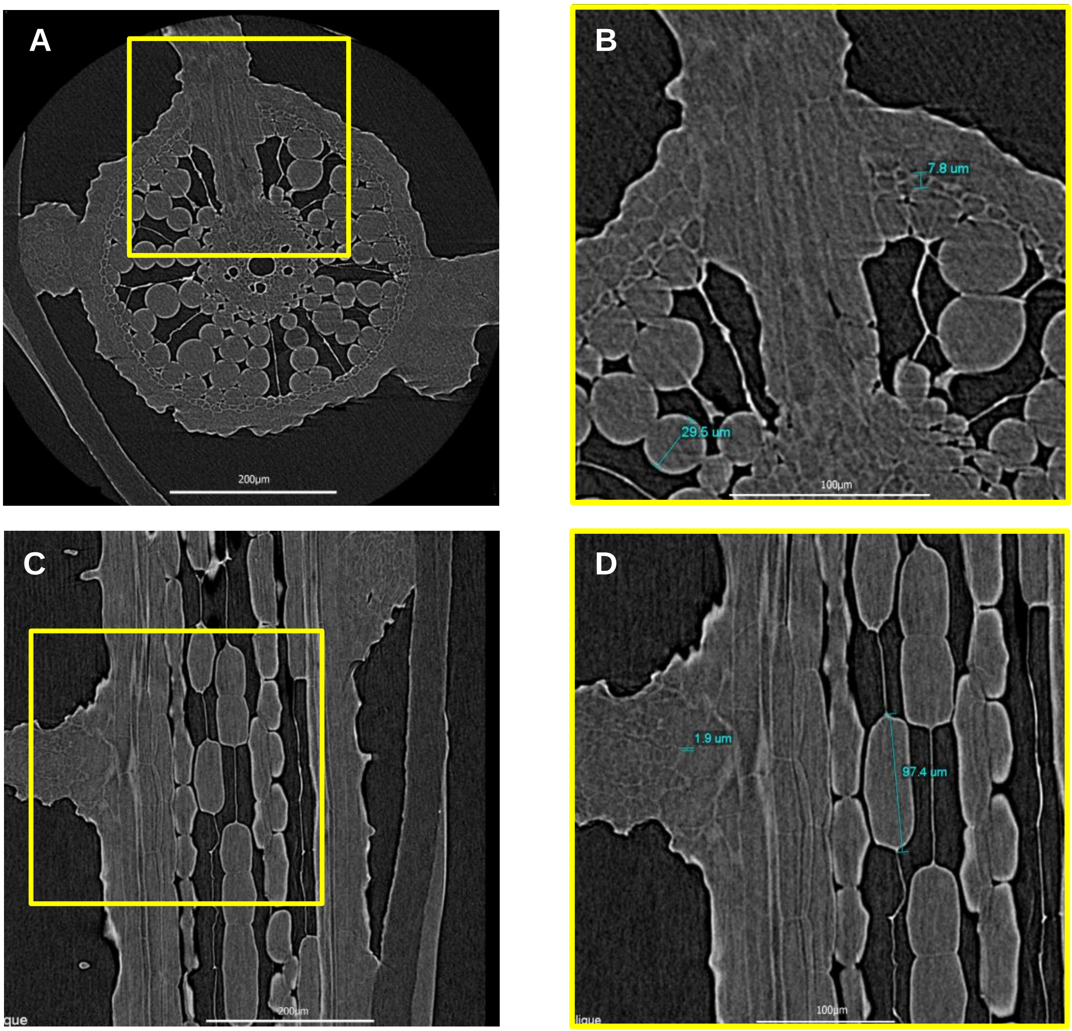


**Figure S1.** Detailed 2D projections of rice root in Fig. 1. (**A**-**B**) 2D projection intersecting the surface of the rice root with 0.7 μm voxel size. Scale bars, 200 μm and 100 μm respectively. Yellow square frame indicates the inset. (**C**-**D**) 2D projection of the lateral surface of the rice root with 0.7 μm voxel size. Scale bars, 200 μm and 100 μm respectively. Individual cell size is measured by blue lines and numbers.

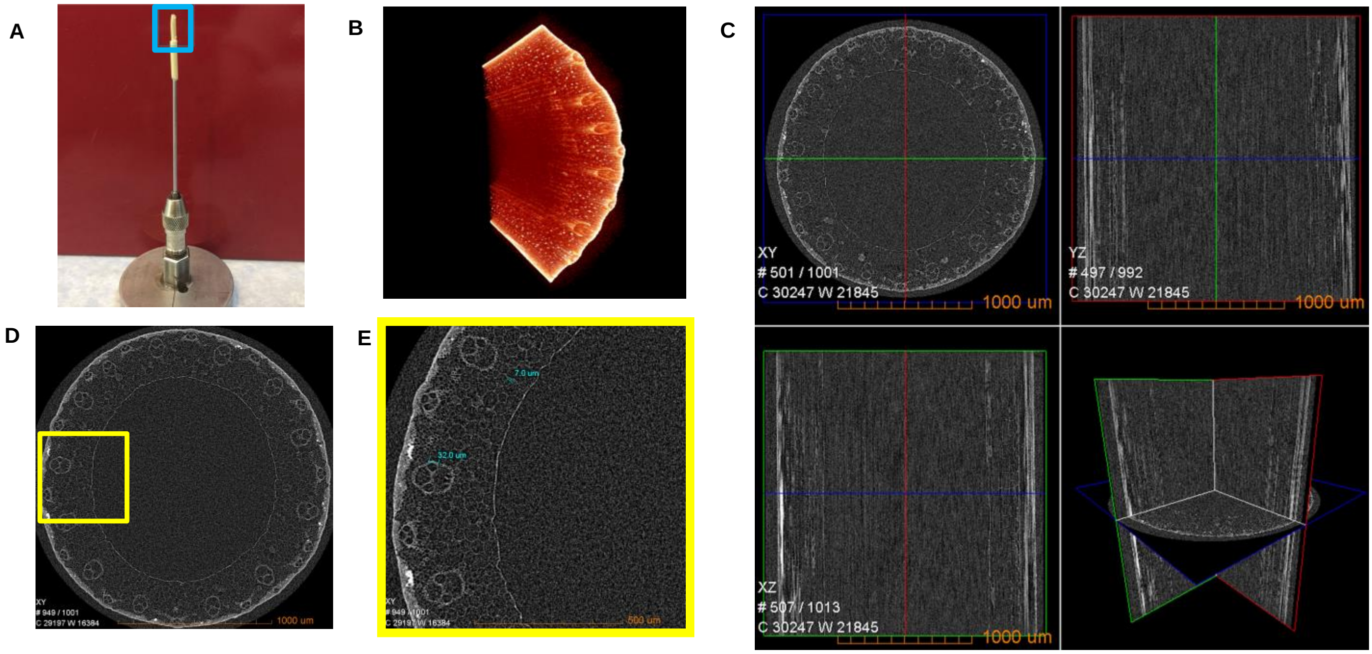


**Figure S2.** The 2.1 μm voxel size scan of the rice stem tissues. (**A**) The loading platform with the mounted rice stem tissue. (**B**) The 3D reconstruction of the sample in (**A**) with computationally rendered. (**C**) 2D projections of GC young stem oriented in three orthogonal directions: the intersecting, longitudinal and lateral cut. Scale bars, 1000 μm. (**D**-**E**) 2D projections intersecting the surface of the stem. Scale bars, 1000 μm and 500 μm respectively.

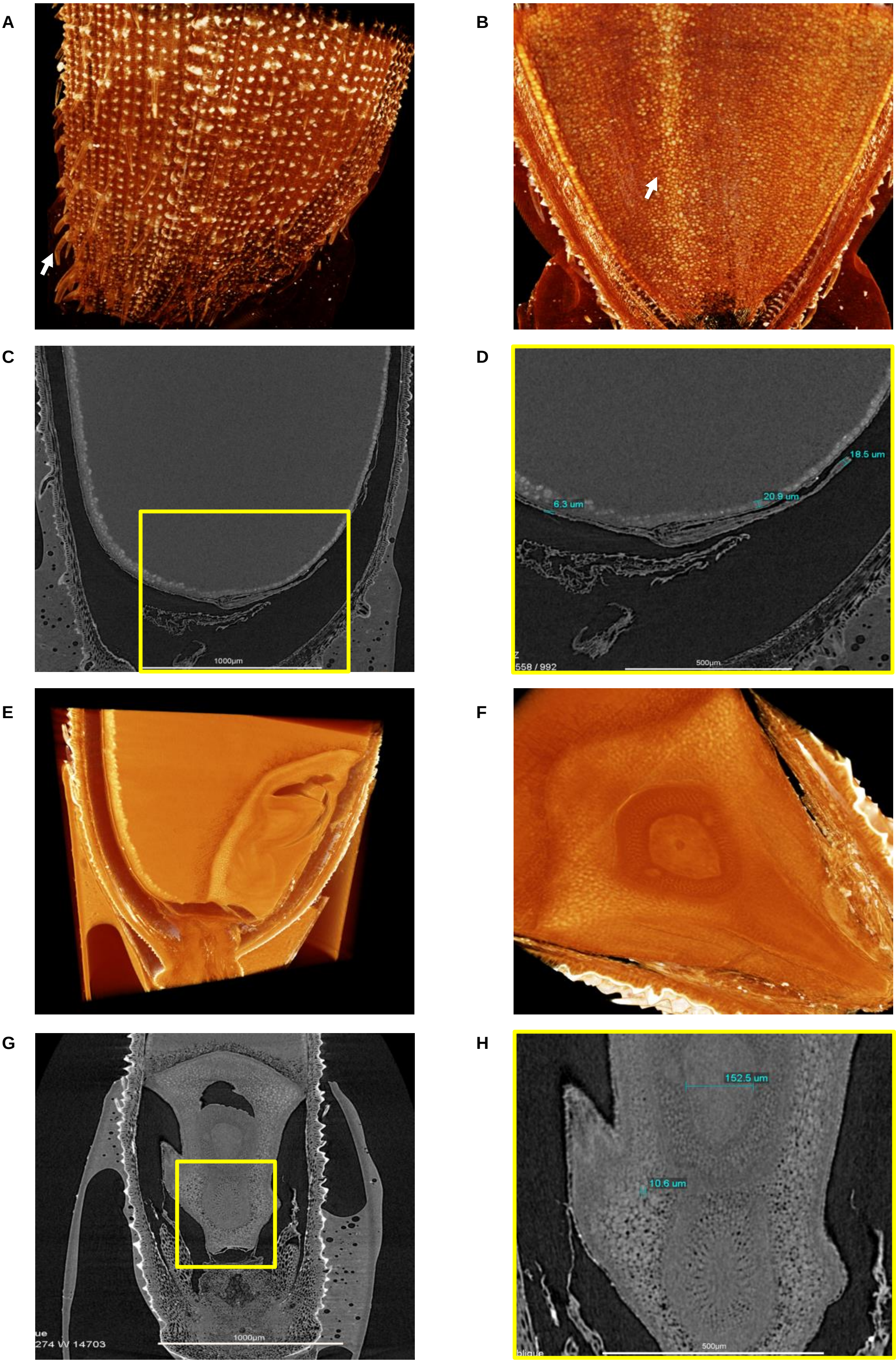


**Figure S3.** Detailed insights into a mature rice grain. (**A**) 3D reconstruction of the rice glume scan at a voxel size of 2.3 μm. The white arrow indicates the glume hair. (**B**) 3D reconstruction of the rice endosperm scan at a voxel size of 2.3 μm. The white arrow indicates the glume hair. The white arrow indicates the starch granules. (**C**-**D**) 2D projection of a longitudinal cut of the top seed with 9.5 μm voxel size. Scale bars, 1000 μm and 500 μm respectively. The yellow square frame indicates the inset. The blue measurements in (**D**) show the different cell widths. (**E**) 3D reconstruction of the rice embryo scan at a voxel size of 2.3 μm. (**F**) The 3D rendering shows the same data as in (**D**) from a more superficial region. (**G**-**H**) 2D projections of a longitudinal cut of the seed embryo with a voxel size of 9.5 μm. Scale bars, 1000 μm and 500 μm respectively. The numbers on (**D**) shows the different tissues width.

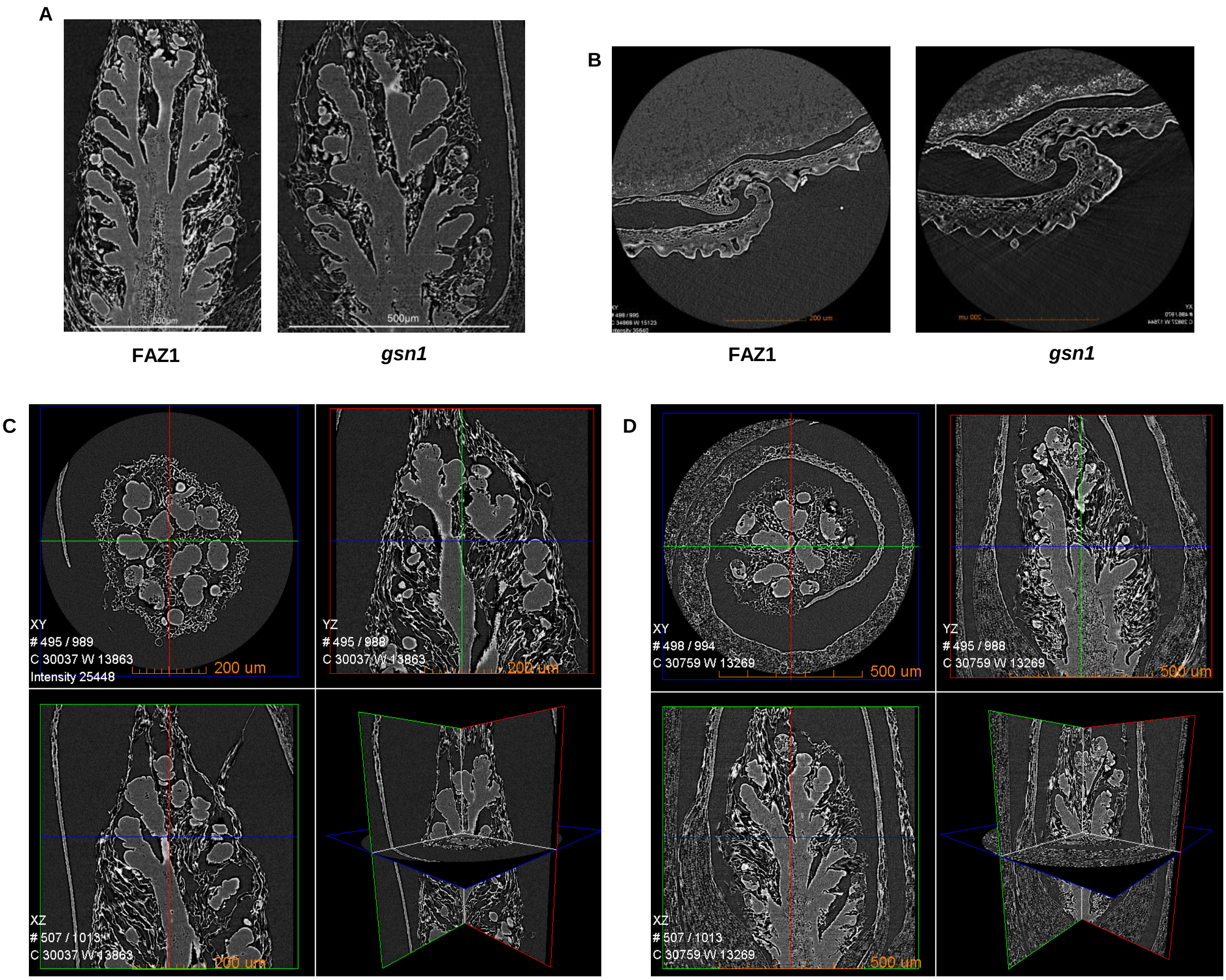


**Figure S4.** Young spikelet and glume comparison of FAZ1 and *gsn1*. (**A**) 2D projection of a longitudinal cut of FAZ1 and *gsn1* young spikelet. Scale bars, 500 µm. The FAZ1 and *gsn1* were scanned with a voxel size of 1.6 µm and 0.9 µm respectively. (**B**) 2D projections intersecting the glumes of FAZ1 and *gsn1*. Scale bars, 200 µm. The FAZ1 and *gsn1* were scanned at a voxel size of 0.7 µm and 0.5 µm respectively. 2D projections of a FAZ1 (**C**) and *gsn1* (**D**) young spikelet in 3 orthogonal orientations: the intersecting, longitudinal and lateral cut. Scale bars, 200 µm. (**D**) 2D projections of young spikelet in 3 different direction. Each direction represents the intersecting cut, longitudinal cut and lateral cut. Scale bars, 500 µm.

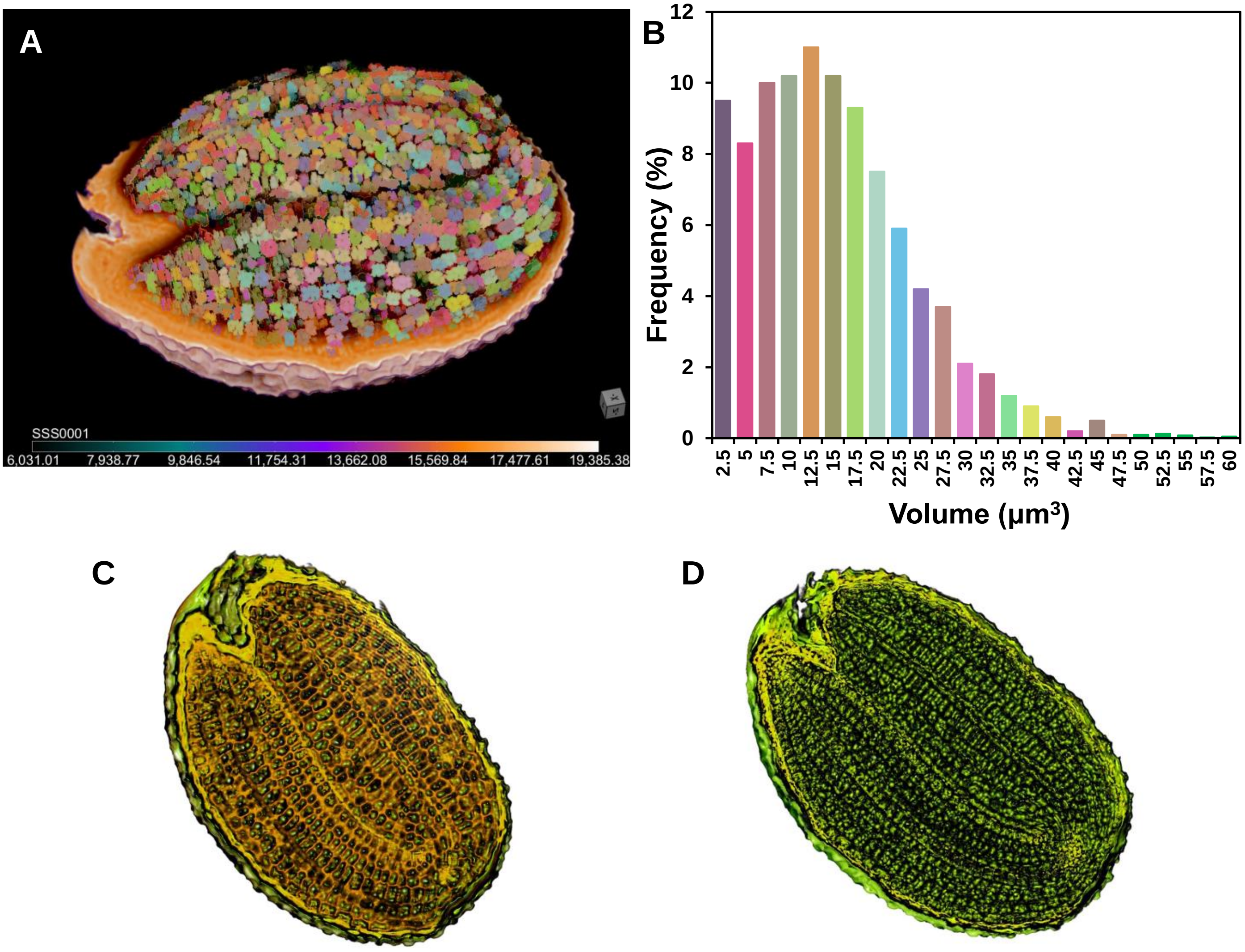


**Figure S5.** Segmentation and rendering of *Arabidopsis* Col-0 seed with the Dragonfly software. (A) The 3D rendering of the Col-0 seed after XRM scanning and data segmentation using the Dragonfly software. Each cell is divided into a separate 3D segmentation and rendered according to the volume. (B) A normalized histogram of the cell volume of the Col-0 seeds. The column colors are consistent with (A). (C-D) Two Col-0 seed 3D renderings color mapped by concavity and convexity (C) and flat embellishment (D).

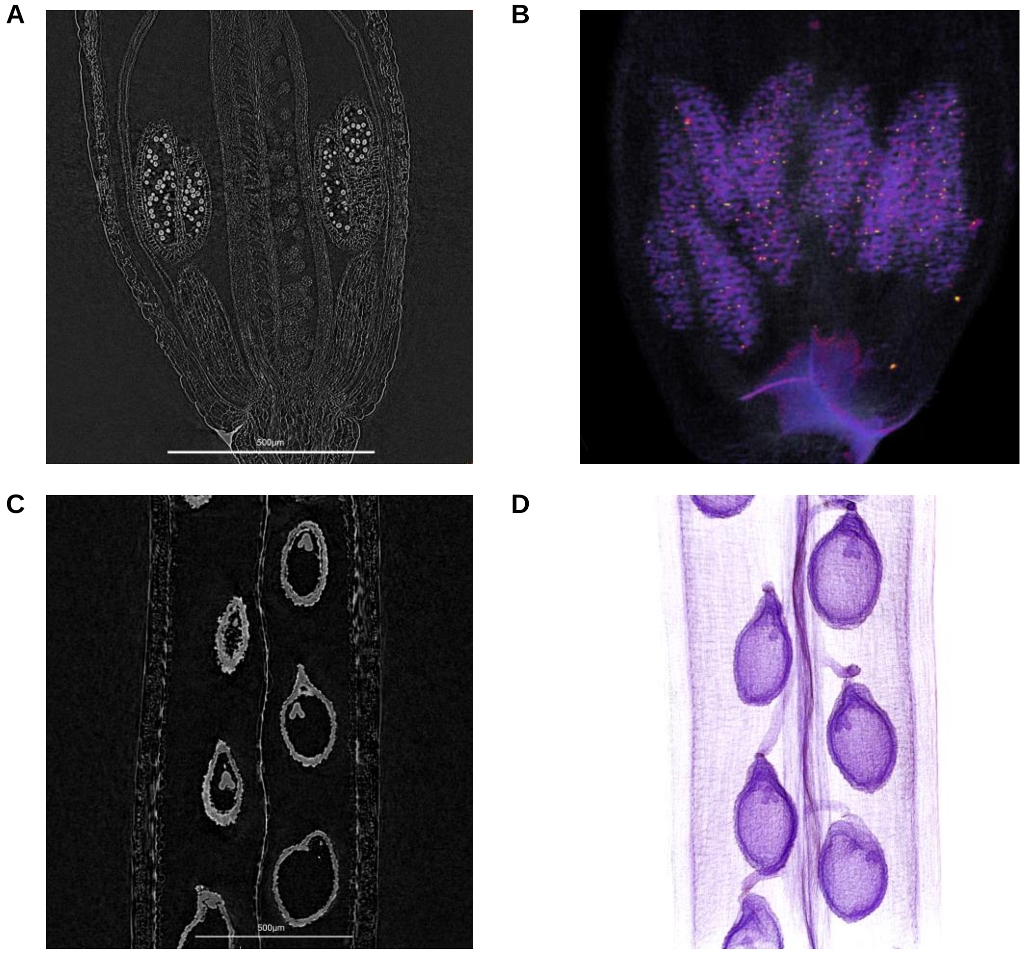


**Figure S6.** The observation of *Arabidopsis* flower buds and legumen. (**A**) 2D projection from a longitudinal cut through the dehydrated Col-0 flower bud. Scale bars, 500 μm. The voxel size of the flower bud scan is 0.9 μm. (**B**) The fake color embellished 3D rendering of the *Arabidopsis* flower bud. Scale bars, 100 μm. (**C**) 2D projection from a longitudinal cut through the Col-0 legumen. Scale bars, 100 μm. (**D**) The pseudo color embellished image of *Arabidopsis* legumen. Scale bars, 500 μm.

### The XRM imaging workflow

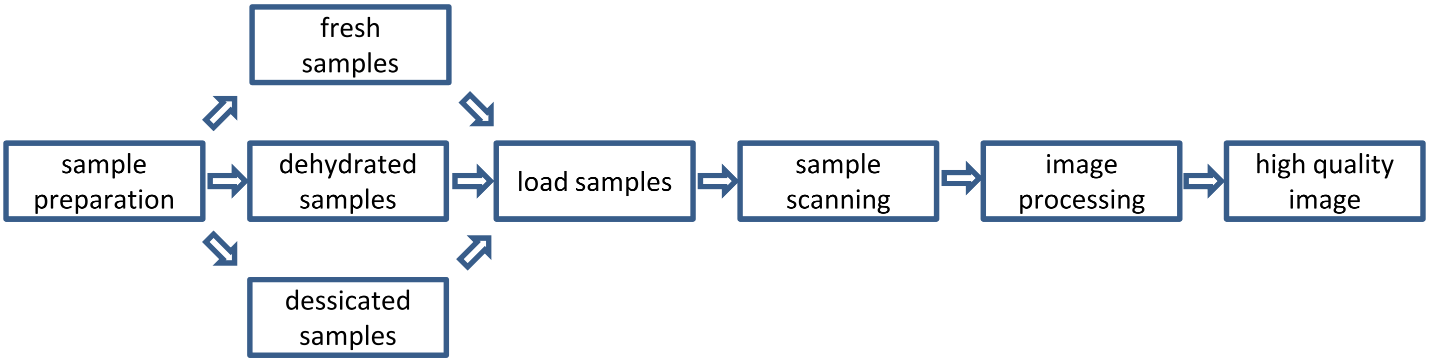


**Figure S7.** The imaging workflow for XRM to observe plant samples. Most plant samples belong to three different categories, including fresh samples, dehydrated samples and desiccated samples. We used CPD method to creat dehydrated samples. After preparation, the samples are loaded in the XRM system. Then, the scanning parameters for each sample are defined and the scan is initiated. During the scanning procedure, the sample should remain still. The scan time depends on the resolution and the x-ray opacity of the sample. The ZEISS system provides powerful and convenient image processing tools to reconstruct the 2D projections.
